# Finding novel vulnerabilities of hypomorphic BRCA1 alleles

**DOI:** 10.1101/2024.05.24.595688

**Authors:** Anne Schreuder, Klaas de Lint, Mariana M. Góis, Rosalie A. Kampen, Marta San Martin Alonso, Ilse Nootenboom, Veronica Garzero, Rob M. F. Wolthuis, Sylvie M. Noordermeer

**Affiliations:** Leiden University Medical Center, Department of Human Genetics, Leiden, The Netherlands; Oncode Institute, Utrecht, The Netherlands; Cancer Center Amsterdam, Amsterdam University Medical Center, Department of Clinical Genetics, Section Oncogenetics, Amsterdam, The Netherlands; State University of Campinas, Laboratory of Applied Molecular Signalling, Campinas, Brazil; Laboratory of Applied Molecular Signalling, Federal University of São Paulo, São Paulo, Brazil

**Keywords:** BRCA1, synthetic lethality, CRISPR Screen, separation of function mutations, BRCA1-R1699Q, NDE1, mitosis

## Abstract

With the recent rise in CRISPR/Cas9-mediated genome-wide synthetic lethality screens, many new synthetic lethal targets have been identified for diseases with underlying genetic causes such as tumours with *BRCA1* mutations. Such screens often use full deficiency of a protein to identify novel vulnerabilities. However, patient-derived mutations not only result in loss of the protein but often also concern missense mutations with hypomorphic phenotypes. Here we study the genetic vulnerabilities of two previously described hypomorphic BRCA1 missense mutations and compare these to a BRCA1-depleted setting to study whether this affects screening for synthetic lethal interactions. Our research showed that BRCA1^I26A^ mutated cells have very similar vulnerabilities to BRCA1 wildtype cells, confirming its low tumorigenic effect. In contrast, the BRCA1^R1699Q^ mutation induced a more similar phenotype to BRCA1-deficient cells. For this mutation, we also unveiled a unique vulnerability to the loss of NDE1. Specifically in BRCA1^R1699Q^ mutated cells, and not BRCA1-proficient or -deficient cells, NDE1 loss leads to increased genomic instability. Altogether our findings highlight the importance to differentiate between patient-derived mutations when assessing novel treatment targets.

## INTRODUCTION

CRISPR/Cas9-mediated genome-wide synthetic lethality screens have shown great potential in unveiling cellular dependency on specific pathways and studying genetic vulnerabilities in certain genomic backgrounds (1-3). Moreover, some of these screens have already resulted in the first clinical trials for certain pathological genetic defects (4) (MYTHIC; Lunresertib; https://doi.org/10.1158/1535-7163.TARG-23-PR008). These screens often use genetically engineered knock-out cell systems to mimic a tumour specific deficiency of the protein of interest. However, such a set-up is not representing missense mutants well, as they often show hypomorphic phenotypes (5). Finding tumour-specific vulnerabilities for patients with specific hypomorphic mutations would thereforecontribute to finding effective treatments for individual patients, in line with the aim of providing personalised medicine.

*BRCA1* is known as a major hereditary breast cancer susceptibility gene, with mutations in *BRCA1* increasing the risk for both breast and ovarian cancer development (6). Moreover, several cancer types also show somatic *BRCA1* mutations and all tumours display a large variety of *BRCA1* mutations. The mutations observed in patients span the full *BRCA1* gene and comprise nonsense, but also missense mutations, many of which result in a protein with hypomorphic function (7). Therefore, the clinical consequence of individual mutations is not always well understood. In addition, the correlation between the type of BRCA1 mutation and their specific treatment response is not always clear. In this paper, we study the genetic vulnerabilities of two previously described hypomorphic *BRCA1* missense mutants and compare these to a BRCA1-depleted setting.

BRCA1 is a key player in the repair of DNA double-stranded breaks (DSBs) via homologous recombination (HR). HR is a very faithful way to repair DSBs since it uses a sister chromatid as an error-free template for repair, limiting HR to S-G2 phase (8,9). During HR, the DNA surrounding the break is resected to expose single-strand DNA (ssDNA). This ssDNA is coated by phosphorylated RPA (pRPA) for protection against nucleases. In the next step, pRPA is replaced by RAD51 filaments which mediates the search for homology and D-loop formation with the sister chromatid allowing for break repair (10-12). BRCA1 stimulates both resection (13,14) and RAD51 loading (10). In addition, BRCA1 is involved in protecting damaged replication forks and prevents ssDNA gap formation (15) and hence BRCA1-defects result in genomic instability.

Studying the BRCA1 c.5096G>A p.Arg1699Gln (R1699Q) variant in the BRCT-domain of BRCA1 has not resulted in a clear conclusion about its functional conseqeunce (16-27) and resulted in an intermediate risk classification (23). BRCA1^R1699Q^ inhibits the binding to the BRCT-binding proteins CTIP, ABRAXAS and BRIP1 (28,29). Individuals carrying this variant show an increased risk of developing breast or ovarian cancer compared to the general population, however the mutation shows reduced penetrance compared to BRCA1 truncating mutations (23,24). BRCA1 is a ubiquitin E3ligase with a RING domain at its N-terminus. With this domain, it forms a dimer with BARD1 which is essential to the stability of the protein. However, the role of the E3 ligase activity for its tumour suppressive function is still under debate (29-32). For example, previous research has shown conflicting effects of the RING I26A mutation, which retains partial binding to BARD1 (29), but disrupts E3 ligase activity (29-34). Although this mutation is not found in patients, it represents a clear example of a hypomorphic BRCA1 mutation.

The conflicting literature on BRCA1^R1699Q^ and BRCA1^I26A^ highlight the uncertainty surrounding the prediction of the functional consequences of hypomophic BRCA1 variants, both for tumour suppression, but also therapy response. In our study, we aim to unveil the genetic vulnerabilities of BRCA1^R1699Q^ and BRCA1^I26A^. To this end, we performed CRISPR/Cas9-mediated genome-wide synthetic lethality screens in RPE1 cells expressing BRCA1^R1699Q^ and BRCA1^I26A^. We have introduced these mutations in a background of endogenously tagged BRCA1-mAID cells (3), in which we can readily deplete BRCA1 using the auxin-inducible degron (AID) (35). This will also allow us to investigate the genetic vulnerabilities of acute BRCA1 loss and compare this to previously reported genetic vulnerabilities of stably generated BRCA1 knock out (KO) cells (1).

Our research reveiled that acute depletion of BRCA1 generates very similar genetic vulnerabilities compared to *BRCA1*-KO cells. In addition, BRCA1^I26A^ has very similar vulnerabilities as BRCA1 wildtype cells, indicating this variant resembles wildtype BRCA1. BRCA1^R1699Q^ induced a more similar phenotype to *BRCA1*-deficient cells, although we also unveiled unique vulnerabilities for this mutation. We found that NDE1, an important protein in centrosome duplication and mitotic spindle formation (36), is important for the survival of BRCA1^R1699Q^ mutated cells specifically. The BRCA1^R1699Q^ variant, in combination with NDE1 loss leads to gross genomic instability and problems during mitosis. The difference in vulnerabilities between BRCA1-depleted and BRCA1^R1699Q^ cells highlights the potential to develop improved tailored personalised treatments based on the precise genetic background of *BRCA1*-mutated tumours.

## RESULTS

### Establishing a model system to test genomic vulnerabilities of the BRCA1 variants BRCA1^R1699Q^ and BRCA1^I26A^

As proof of concept, we set out to compare BRCA1-depleted cells with BRCA1-mutated R1699Q and I26A cells. To enable our studies on these variants, we used our previously generated RPE1 hTERT *TP53*^-/-^ cell line containing an auxin-inducible degron (mAID) fused to the endogenous *BRCA1* gene (3,35) and virally complemented these cells with doxycycline inducible Ty1-tagged BRCA1 wildtype, BRCA1^R1699Q^ or BRCA1^I26A^. Treatment of these cells with auxin readily depleted the endogenous levels of BRCA1 as described before (3) and doxycycline induced the expression of the BRCA1 variants (Supplemental figure 1A) to levels comparable to the endogenous protein.

BRCA1^R1699Q^ has been described to inhibit the binding to the BRCT-binding proteins CTIP, ABRAXAS and BRIP1 (28,29). An immunoprecipitation using antibodies against the Ty1 tag of BRCA1 validated that this mutation disrupts binding to CTIP, ABRAXAS and BRIP1 (Supplemental figure 1B). Next, we examined whether the two mutations affected BRCA1’s recruitment to DSBs by analysing ionising irradiation-induced foci (IRIF) formation. In addition, we analysed RAD51 IRIF as a read-out for HR. As expected, after depletion of BRCA1 both BRCA1 and RAD51 recruitment was impaired and this was rescued by re-expression of exogenous wildtype BRCA1 (Figure 1A). BRCA1^R1699Q^ showed a major decrease in BRCA1 IRIF compared to the wildtype, as was described previously for this BRCA1 variant (19). In line with the reduced BRCA1 IRIF, less RAD51 foci were formed in the BRCA1^R1699Q^ variant (Figure 1B). BRCA1^I26A^ also negatively affected BRCA1 IRIF compared to wildtype (Figure 1A). Even though the reduction in BRCA1 IRIF was not as severe as for BRCA1^R1699Q^, RAD51 IRIF were reduced to similar levels for both mutants (Figure 1B).

**Figure 1.**
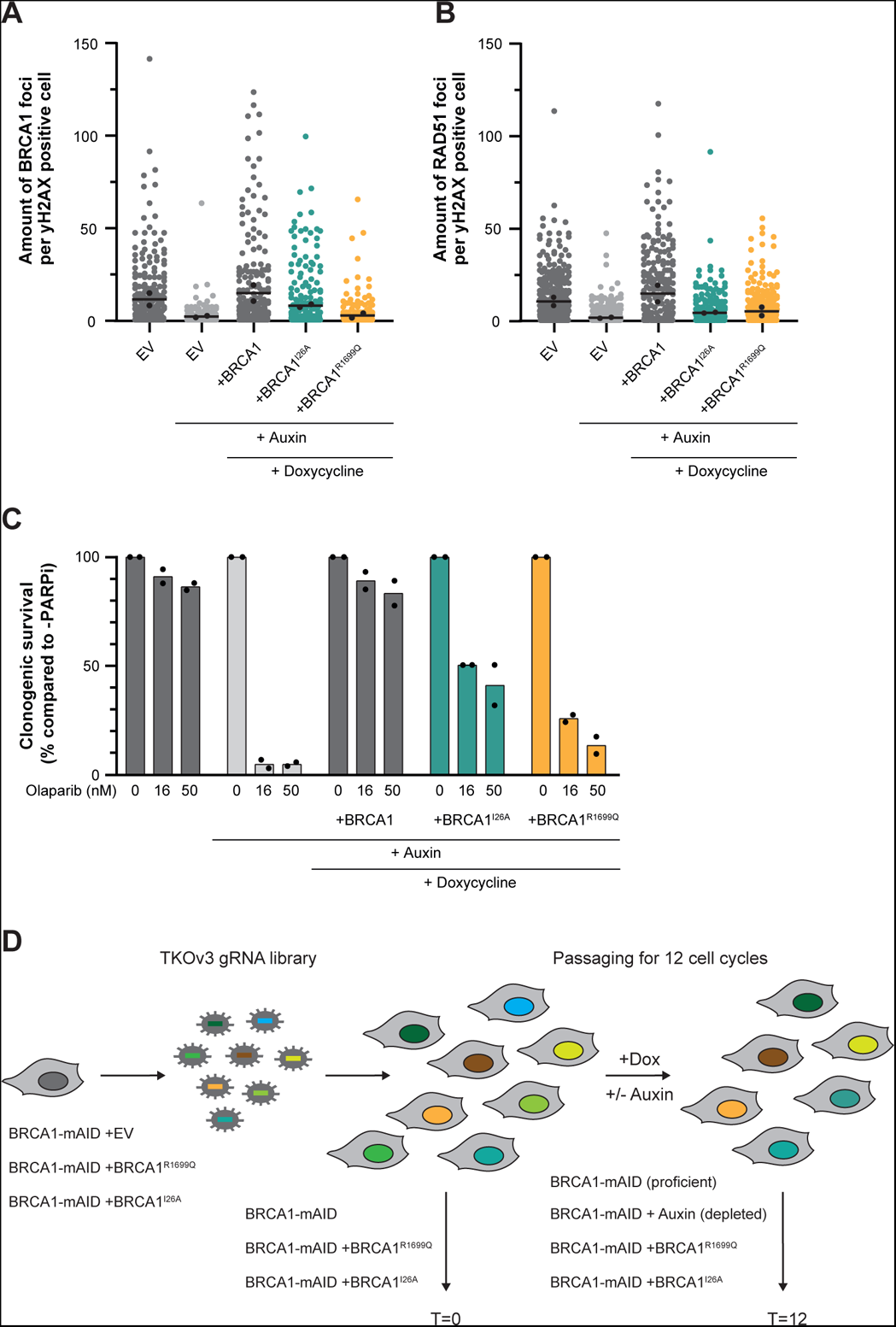
Validation of the BRCA1-mutant phenotypes. (A) The indicated complemented RPE1 hTERT *TP53*^-/-^ *BRCA1-mAID* cell lines were irradiated with 10 Gy and subsequent IF microscopy was performed to analyse BRCA1 foci formation 3 hrs post-IR (n=2, mean). Western blot of the used cells is shown in Supplemental figure 1A. **(B)** The indicated complemented RPE1 hTERT *TP53*^-/-^ *BRCA1-mAID* cell lines were irradiated with 10 Gy and subsequent IF microscopy was performed to analyse RAD51 foci formation 3 hrs post-IR (n=2, mean). Western blot of the used cells is shown in Supple-mental figure 1A. **(C)** The indicated complemented RPE1 hTERT *TP53*^-/-^ *BRCA1-mAID* cell lines were treated with different concentrations of the PARPi Olaparib and viability was assessed using a clonogenic survival assay. (n=2, mean). Western blot of the used cells is shown in Supplemental figure 1A. **(D)** Graphical representation of the genome-wide CRISPR-Cas9 knockout screen set-up. The four conditions in the screen were: 1) BRCA1-mAID (BRCA1-proficient), 2) BRCA1-mAID with 500 µM auxin treatment which readily led to the depletion of BRCA1, 3) BRCA1-mAID +BRCA1^R1699Q^ with auxin treatment to deplete endogenous BRCA1 and treatment with Doxycycline to express BRCA1^R1699Q^ and 4) BRCA1-mAID +BRCA1^I26A^ with auxin treatment to deplete endogenous BRCA1 and treatment with Doxycycline to express BRCA1^I26A^. The screen was performed in triplicate for each condition.

To assess the cellular consequence of these mutations, we performed clonogenic survival assays with the PARP inhibitor (PARPi) Olaparib (37). As expected, Olaparib treatment led to cell death in BRCA1-depleted cells (Figure 1C). Re-expression of wildtype BRCA1 led to a full rescue of the sensitivity to Olaparib, whereas BRCA1^I26A^ only partially rescued the sensitivity (Figure 1C). Although severe Olaparib sensitivity of BRCA1^I26A^ has been described previously (29), our results are more in line with the described capacity of BRCA1^I26A^ cells to perform HR (29,30). BRCA1^R1699Q^ showed a more severe sensitivity to Olaparib as was described previously in mESCs (38) and MDA-MB-436 breast cancer cells (27,29), although it was more resistant than BRCA1-depleted cells. Altogether our data confirm that both variants show a hypomorphic phenotype with the BRCA1^R1699Q^ variant showing a more severe phenotype than the BRCA1^I26A^ variant.

### Setting up genome-wide CRISPR-Cas9 knockout screens to assess BRCA1-mutant vulnerabilities

To unveil the specific vulnerabilities of the different BRCA1 variants, we performed a CRISPR/Cas9-mediated genome-wide synthetic lethality screen in RPE1 cells expressing BRCA1^R1699Q^ and BRCA1^I26A^. As preparation for the screen, we virally integrated an inducible Cas9 casette (Supplemental figure 1A) into our RPE1 hTERT *TP53*^-/-^ *BRCA1-mAID* cells. We confirmed its functionality by observing > 95% cell death upon expression of doxycycline-induced Cas9 combined with the overexpression of a sgRNA against the essential gene *PSMD1* (Supplemental figure 1C). Subsequently, the inducible Cas9 cell clone was virally complemented with the different BRCA1 variants.

To identify synthetic lethal interactions for the specific BRCA1 variants, we included four arms in our genome-wide CRISPR-screen: 1) *BRCA1-mAID* (*BRCA1*-proficient; no auxin), 2) *BRCA1-mAID* with 500 µM auxin treatment (*BRCA1*-depleted), 3) BRCA1-mAID +BRCA1^R1699Q^ with auxin and 4) BRCA1-mAID +BRCA1^I26A^ with auxin (Figure 1D). All populations were cultured in the presence of doxycycline to induce Cas9 and BRCA1 variant (for arm 3-4) expression. For each condition, the cells were transduced with the TKOv3 sgRNA library and grown for 12 doublings before collection of genomic DNA. As the doubling time of the different cell lines differed – with the BRCA1-depleted and BRCA1^R1699Q^ mutant showing the slowest cell growth (Supplemental figure 1D) – we adjusted the time to reach 12 doubling per condition.

*Screening for essential genes upon acute BRCA1-loss resembles screens of clonal BRCA1-deficient cells* Our set-up of using auxin-inducible BRCA1 depletion allowed us to study the genetic vulnerabilities of BRCA1 loss in cells that did not adapt to BRCA1 loss over time (as might be the case with full *BRCA1*-deficient cells). Genes known to have a synthetic lethal interaction with BRCA1 loss, such as *APEX2*, *PARP1* and *POLQ* were also strong hits in our CRISPR screen comparing BRCA1-proficient and -depleted cells (Figure 2A, Supplemental table 1) (39-42). Furthermore, genes that are known to stimulate survival of *BRCA1*-deficient cells when defective, like *TP53BP1* and the Shieldin complex, were stimulating survival of BRCA1-depleted cells in our CRISPR screen as well (2,43-48). This suggests that our inducible BRCA1 depletion results in a cellular phenotype resembling *BRCA1* deficiency.

**Figure 2.**
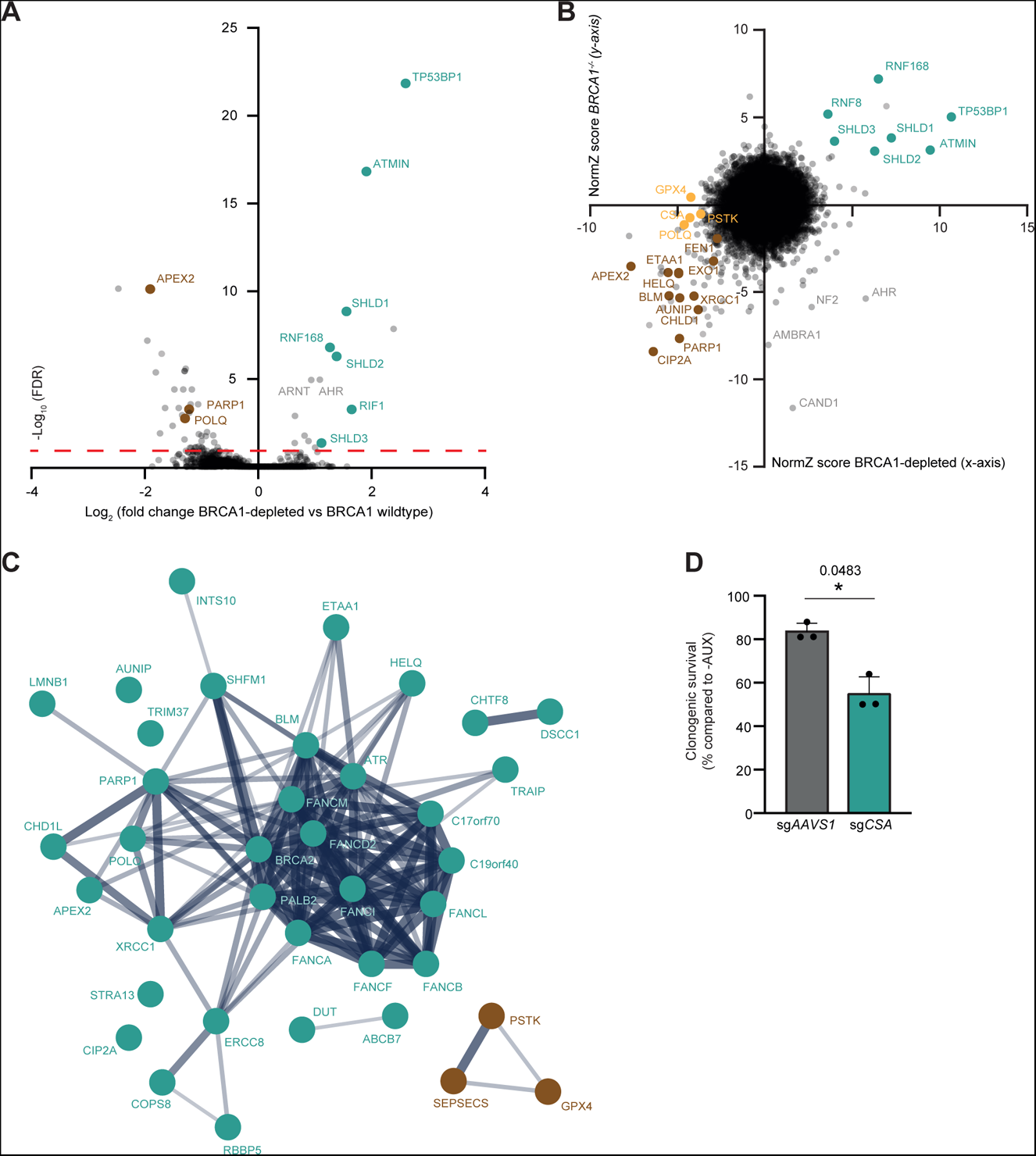
Screening for essential genes upon acute BRCA1-loss resembles screen of clonal BRCA1-deficient cells. (A) Volcano plot depicting genes that are synthetic lethal with BRCA1 depletion (left side) or promote survival of cells that suffered acute BRCA1-loss (right side, green data points). Brown data points indicate previously described synthetic lethal interactions with BRCA1-deficiency. The fold change (Log2) is plotted on the x-axis and the significance (-Log10 false discovery rate) on the y-axis with the cut-off line in red at -Log10(0.1)=1. **(B)** The NormZ score of each gene in the BRCA1-depleted (TKOv3) screen and the reanalysed *BRCA1^-/-^* screen (TKOv2, Adam et al. 2021) were plotted. 156 genes were in the TKOv3 but not TKOv2 version of the sgRNA library, 45 genes were in TKOv2 but not TKOv3, leaving 17,900 pairs for the correlation plot (n=17,900). The coloured data points indicate previously described synthetic lethal or rescue genes when lost in a BRCA1-deficient background. **(C)** STRING pathway analysis for all hits (FDR< 0.1) that are synthetic lethal with BRCA1-depletion. The line thinkness indicates the strength of the data support from textmining, experiments and databases. **(D)** RPE1 hTERT *TP53*^-/-^ *BRCA1-mAID* cell lines were virally transduced to express *Cas9* cDNA and a sgRNA against *AAVS1* or *CSA*, followed by a clonogenic survival assay in presence or absence of 500 μM Auxin (n=3, mean+SD, *p<0.05, ratio paired t-test)). Western blot of lysates shown in Supplemental figure 2A.

For a direct comparison between acute BRCA1-loss and clonal *BRCA1*-deficient cells, we compared our screen results with a previously performed CRISPR screen in *BRCA1*-deficient RPE1 cells by Adam *et al.* (1). For this, we re-analysed the CRISPR screen in *BRCA1*-deficient cells using IsogenicZ, an adaptation of DrugZ, which we used to analyse our screens (https://github.com/kdelint/IsogenicZ) (1). Based on the correlation plot between the two screens it is clear that there is strong overlap between the two screens as the majority of positive and negative hits are shared in the two screens (Figure 2B). Also the genes *CIP2A*, *ETAA1* and *AUNIP*, first identified by Adam *et al.* were synthetic lethal hits for the BRCA1-depleted cells in our screen (1). A striking difference was the known SL interactor *POLQ* (Figure 2B) being a hit in the case of acute BRCA1-loss and not in clonal *BRCA1*-deficient cells. However, overall there were only few differential hits between the BRCA1-depleted and -deficient screen, confirming that adaptation of *BRCA1*-deficient cells is not a major drawback in this case.

Pathway analysis shows that the majority of the synthetic lethal hits for the BRCA1-depleted cells are related to DNA repair with an FDR of 4.17e^-25^ using Gene Ontology analysis on Biological processes and an FDR of 5.93e^-21^ using Reactome Pathway analysis (Figure 2C). One hit related to DNA repair, but not previously described as synthetic lethal with BRCA1 loss, is CSA (ERCC8). CSA is an important protein in DNA repair via transcription-coupled nucleotide excision repair (TC-NER) (Figure 2C). We validated this synthetic lethal interaction by dedicated experiments in BRCA1-depleted RPE1 cells and BRCA1-mutated HCC1937 cells (Figure 2D, Supplemental figure 2A, B, C).

In addition to genes linked to DNA repair, we found glutathione peroxidative 4 (*GPX4*) and phosphoseryl-tRNA kinase (*PSTK*) loss as synthetic lethal in BRCA1-depleted cells (Figure 2B, C). Both proteins are important for the protection of the cell against ferroptosis (49,50) and loss of GPX4 has been described to abolish tumour growth and enhance treatment sensitivity (51-54). Moreover, GPX4 inhibition has recently been described to induce cell death in *BRCA1*-deficient cells (55). However, we would like to add a word of caution here, as there is also clear BRCA1-status independent toxicity upon GPX4 loss when checking the raw data of the screen (data not shown). Indeed, also in the previous report, the authors show toxicity upon GPX4 inhibition in *BRCA1*-proficient cells depending on the cell line used (55).

### BRCA1 ^I26A^ exhibits vulnerabilities similar to BRCA1-proficient cells

When comparing the essential genes identified in the BRCA1^I26A^ and BRCA1^R1699Q^ cells to the hits found in the BRCA1-depleted background, it is clear that the BRCA1^I26A^ mutant only shows little synthetic lethality hits, as summarised by the Venn diagram in Figure 3A. The majority of the BRCA1^I26A^ hits (*FANCM*, *FAAP24 (C19orf40)*, *FANCD2*) are Fanconi Anemia (FA) core complex proteins and among the strongest hits for BRCA1-depleted cells as well. The lack of other significant differences between BRCA1^I26A^ and *BRCA1*-proficient cells highlights the low impact of this mutation on BRCA1 function (Figure 3B, Supplemental table 2). This finding correlates with research showing that the I26A mutation does not affect tumour suppression and HR by BRCA1 (30,31). All together, this exemplifies that the BRCA1^I26A^ variant has a mild phenotype and shows high similarity to *BRCA1*-proficient cells.

**Figure 3.**
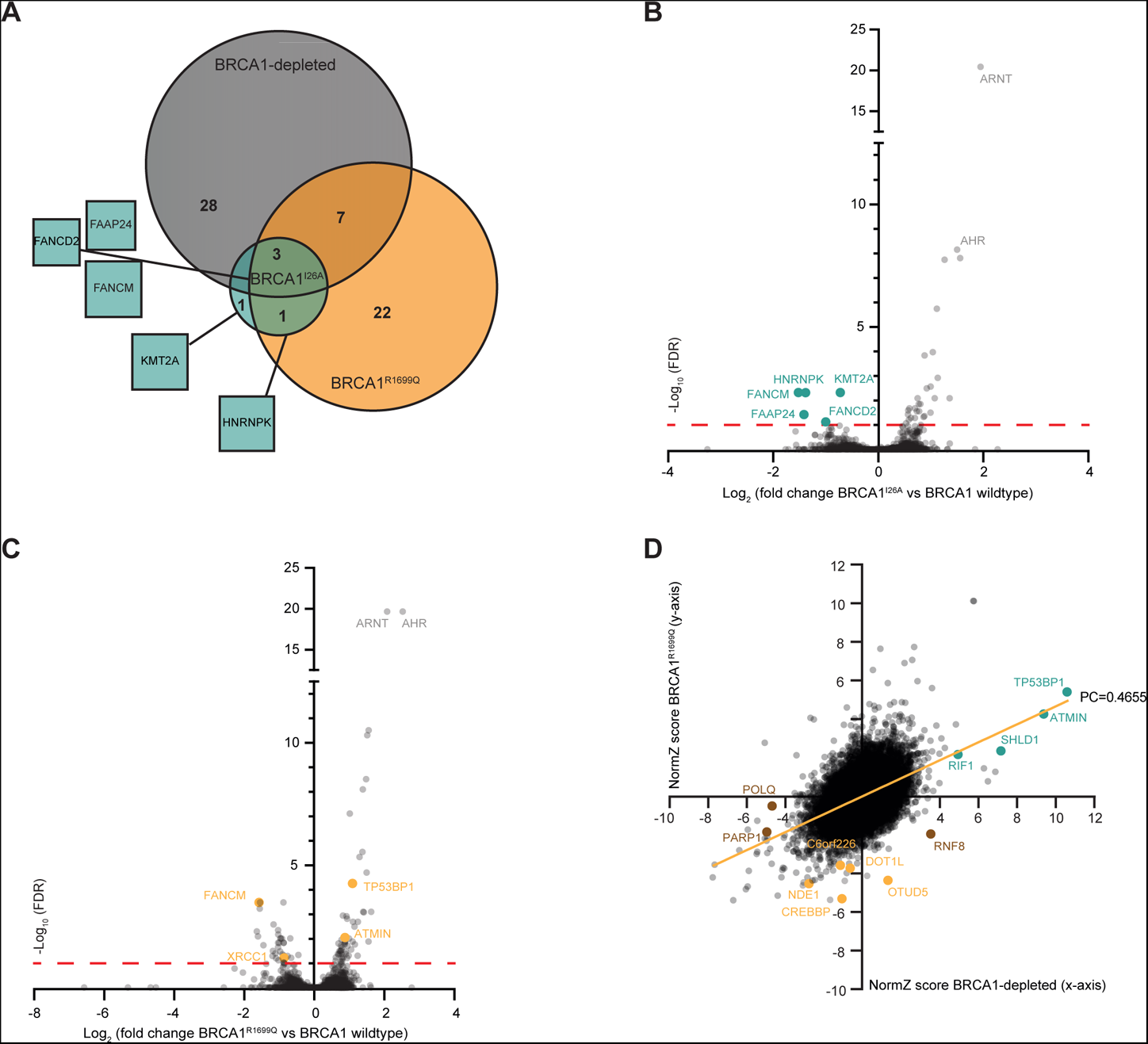
Genetic vulnerabilities and resistances differ between BRCA1^R1699Q^ and BRCA1-depleted cells. (A) Venn diagram comparing the different arms of the genome-wide CRISPR-Cas9 knockout screen. The hits of the represented arms are compared to the BRCA1-proficient arm of the screen. The BRCA1^I26A^ hits are shown by name, the square size repesents the BRCA1^I26A^vsBRCA1-proficient NormZ score of the respective hit. **(B)** Volcano plot depicting genes that are synthetic lethal with BRCA1^I26A^ (left side) or promote survival of cells with BRCA1^I26A^ (right side). The fold change (Log2) is plotted on the x-axis and the significance (-Log10 false discovery rate) on the y-axis with the cut-off line in red at -Log10(0.1)=1. **(C)** Volcano plot depicting genes that are synthetic lethal with BRCA1^R1699Q^ (left side) or promote survival of cells with BRCA1^R1699Q^ (right side). The fold change (Log2) is plotted on the x-axis and the significance (-Log10 false discov-ery rate) on the y-axis with the cut-off line in red at -Log10(0.1)=1. **(D)** NormZ score of each gene in the BRCA1-depleted screen and the NormZ score of each gene in the BRCA1^R1699Q^ screen were plotted (n=18,055). Orange line is the linear regression curve, PC=Pearson correlation coefficient. The coloured data points indicate interesting genes.

### Unique vulnerabilities for cells expressing BRCA1^R1699Q^

The BRCA1^R1699Q^ variant screen resulted in many more hits than the BRCA1^I26A^ screen (Figure 3A) and many hits overlap with the hits found in the BRCA1-depleted cells, such as *XRCC1* and several FA genes (Figure 3C, Supplemental table 3). All three BRCA1-statuses showed loss of AHR or ARNT is beneficial for survival in the screen, which could be due to an effect of auxin (56). Furthermore, genes that are known to rescue survival-defects of *BRCA1*-deficient cells, like *TP53BP1* and *ATMIN*, were also rescuing the BRCA1^R1699Q^ cells in our CRISPR screen (Figure 3C, D). However, the genetic vulnerabilities and resistances found for BRCA1^R1699Q^ cells and BRCA1-depleted cells, do show some dissimilarities (Figure 3A, D). For example, the loss of *RNF8* stimulates proliferation of *BRCA1*-deficient cells but not of BRCA1^R1699Q^ cells, which we confirmed in a dedicated cellular growth assay (Figure 3D, Supplemental figure 2D).

Other examples of differences between *BRCA1*-deficient and BRCA1^R1699Q^ cells are *PARP1* and *POLQ*, which are, unexpectedly, no strong synthetic lethal targets for BRCA1^R1699Q^ cells (Figure 3D). Currently, PARPi are used in the clinic to treat patients with BRCA1 mutations and for POLQ inhibitors clinical phase II research has started (CT169; GSK4524101; https://doi.org/10.1158/15387445.AM2024-CT169, ART4215; Artios). Interestingly, both PARP1 and POLQ loss are not very toxic to the BRCA1^R1699Q^ mutated cell in our CRISPR screen. Likewise, in the clonogenic survival assay, the BRCA1^R1699Q^ variant was also less sensitive to Olaparib compared to the BRCA1-depleted cells (Figure 1C). In contrast, ClinVar (57) categorises BRCA1^R1699Q^ as pathogenic and therefore patients with this mutation are likely treated with PARPi. Our data suggest that this current treatment might not be fully effective for tumours with such BRCA1 mutation.

### Loss of NDE1 is specifically toxic to cells expressing BRCA1^R1699Q^

Pathway analysis on the top vulnerabilities (NormZ < -2.5) for BRCA1^R1699Q^ cells revealed a strong enrichment of proteins associated with mitotic spindle assembly and checkpoint, an association less clear for the BRCA1-depleted hits (Figure 4A). This was especially true for six specific processes, namely mitotic spindle checkpoint, separation of sister chromatids, amplification of signal from the kinetochores, amplification of signal from unattached kinetochores via a MAD2 inhibitory signal, resolution of sister chromatid cohesion, EML4 and NUDC in mitotic spindle formation (Supplemental figure 3A). The genes *NDE1*, *PPP2CA* and *DYNC1LI1* were a significant hit (FDR < 0.1) in the screen for BRCA1^R1699Q^ cells compared to *BRCA1*-proficient cells (Figure 4A) and were involved in all six of these processes. The other significant hits *CDCA5* and *PSMD13* however, are only involved in one or two of the specific processes. Based on the NormZ score, we followed up on NDE1, an essential protein in spindle assembly (see below).

**Figure 4.**
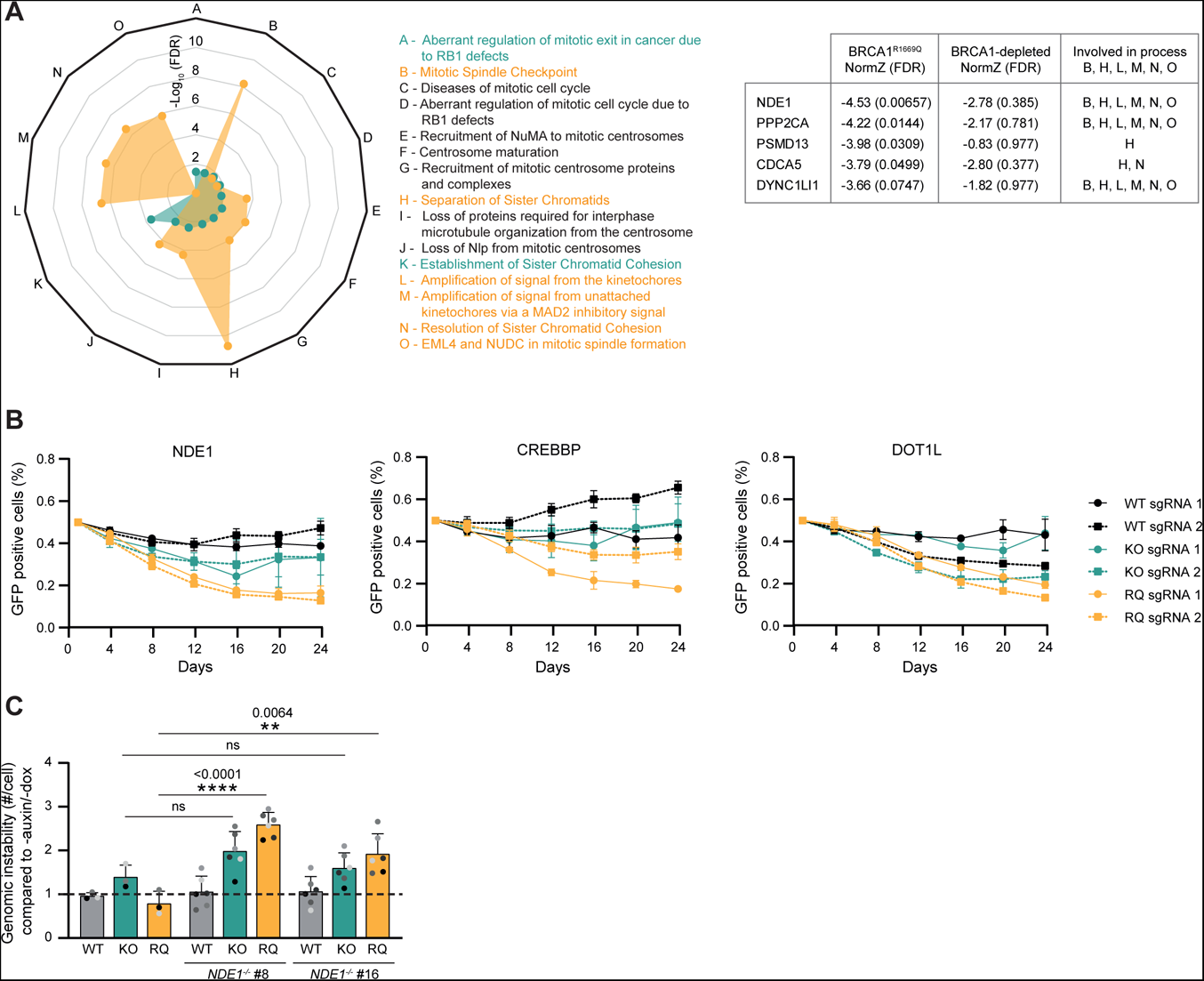
NDE1 loss is toxic for cells with BRCA1^R1699Q^. (A) Panther pathway analysis on the top vulnerabilities (NormZ < -2.5) of BRCA1-depleted cells (green) and BRCA1^R1699Q^ cells (orange). Pathways enriched with an FDR < 0.05 were considered statistically significant. The identified pathways per arm were manually curated for pathways involved in mitosis and compared between the two arms.The significance of the involvement of the vulnerabilities in each mitotic process is plotted (-Log10 false discovery rate). Supplemental figure 3A shows the raw number of genes involved in the specific mitotic process for the two different BRCA1 statuses. **(B)** RPE1 BRCA1-mAID cells expressing doxycyclin inducible Cas9, either *BRCA1^+/+^* (WT, black lines), BRCA1-depleted with auxin (KO, green lines) or BRCA1^R1699Q^ complemented (RQ, orange lines), were infected with indicated sgRNA together with GFP, or with an empty vector together with mCherry. GFP- and mCherry-positive cells were mixed 1:1, and the ratio of GFP-positive cells in the population was determined over time (n=3, mean±SEM). gRNA efficiency is shown in Supplemental figure 3C. **(C)** The indicated complemented (with either EV, doxy-cycline inducible BRCA1-WT or BRCA1^R1699Q^) RPE1 hTERT *TP53*^-/-^ *BRCA1-mAID NDE1^+/+^* or *NDE1^-/-^* cell lines were grown with or without auxin (to deplete endogenous BRCA1) and doxycycline (to express the BRCA1 complementation) for 48 hours before fixation. Genomic instability is the average amount of micronuclei (Supplemental figure 3E) and anaphase bridges (Supplemental figure 3F) per cell. (n=3 and n=6, mean+SD, ns p>0.05, **p<0.01, ****p<0.0001, unpaired t-test) Western blot of the used cells is shown in Supplemental figure 3D.

Additionally, we included other strong unique hits specifical for BRCA1^R1699Q^ in our validation assays (Figure 3D): *CREBBP, DOT1L, OTUD5* and *C6orf226*. To validate the differential BRCA1^R1699Q^ hits, we performed a MCA as described previously (2). In short, cells were transduced with lentivirus either expressing an sgRNA targeting a gene-of-interest and GFP or mCherry only. These populations were mixed 1:1 and the ratio of mCherry-positive and GFP-positive cells was monitored over time. Confirming data from the CRISPR screen, depletion of NDE1, CREBBP, DOT1L or OTUD5 led to enhanced toxicity in BRCA1^R1699Q^ expressing cells compared to BRCA1-depleted or -proficient cells (Figure 4B, Supplemental figure 3B). We could not validate the toxicity of C6orf226 loss, however this could be due to poor gene editing efficiency (Supplemental figure 3B, C).

To further investigate the synthetic lethal interaction of NDE1 loss in BRCA1^R1699Q^ cells, we generated RPE1 hTERT *TP53^-/-^ BRCA1-mAID NDE1^-/-^* cells which we complemented with an empty vector plasmid, BRCA1 wildtype or BRCA1^R1699Q^ (Supplemental figure 3D). NDE1 is important for centrosome duplication, mitotic spindle formation and loss of NDE1 has been found to increase DSBs (36,58). Besides the role of BRCA1 in DSB repair, replication fork protection and ssDNA gap prevention, the protein has also been linked to mitosis and more specifically to the regulation of centrosome duplication (59,60). Since both BRCA1 and NDE1 play an important role in processes maintaining genomic stability, we studied genomic instability in our different cell lines. BRCA1-depleted cells, independent of NDE1 status, show more genomic instability in the form of micronuclei and anaphase bridges compared to *BRCA1* proficient cells as expected (Figure 4C). Interestingly, in a BRCA1^R1699Q^ background, only *NDE1^-/-^* cells and not *NDE1^+/+^* show more genomic instability compared to BRCA1 proficient and even BRCA1-depleted cells (Figure 4C, Supplemental figure 3E,F). All together, the increased genomic instability of NDE1 loss in BRCA1^R1699Q^ cells could explain the poor survival of these cells.

## DISCUSSION

CRISPR screens have great potential to identify novel targets for specialised treatment of diseases caused by genetic defects. Therefore, there has been a surge in screens using knock-out settings to find novel synthetic lethal interactions. However, in many diseases, there is large heterogeneity in missense mutations between patients and not all mutations lead to a complete loss-of-function of the protein. In this study, we therefore compared synthetic lethal interactions of two hypomorphic mutations in BRCA1 - BRCA1^I26A^ and BRCA1^R1669Q^ - to a BRCA1-depleted background. In the phenotypical assays, analysing BRCA1 IRIF and PARPi sensitivity, the BRCA1^I26A^ mutant cells showed a mild phenotype and the BRCA1^R1699Q^ cells exhibited a more severe phenotype. This also correlated to the screen results, where BRCA1^R1699Q^ mutated cells showed more vulnerabilities compared to BRCA1^I26A^ mutated cells.

In our analysis of the vulnerabilities of BRCA1-depleted cells we uncovered a hitherto unknown synthetic lethal interaction between CSA and BRCA1 loss. Previously, it has been suggested that BRCA1 and CSA can independently polyubiquitinate CSB, a process important during TC-NER (61). This suggests that BRCA1 might play an auxiliary role to CSA which could explain the synthetic lethal interaction between BRCA1 and CSA.

Our results highlight the mild phenotype of BRCA1^I26A^ mutated cells. This correlates to research showing that the I26A mutation does not affect tumour suppression and HR (26,27). Furthermore, recently it has been found that full-length BRCA1^I26A^-BARD1 retains substantial ubiquitin E3 ligase activity and that a triple BRCA1 variant (I26A/L63A/K65A) is needed to fully disrupt ligase activity (62). Together, these results indicate that the BRCA1^I26A^ mutation has low impact on BRCA1 function.

While the BRCA1^R1699Q^ mutated cells resembled BRCA1-depleted cells more closely, we found specific vulnerabilities or resistances that were distinct between BRCA1-depleted cells and BRCA1^R1699Q^ mutated cells. For example, loss of *RNF8* stimulates proliferation of BRCA1-depleted cells but not BRCA1^R1699Q^ or *BRCA1*-proficient cells. Loss of *RNF8* protecting *BRCA1*-deficient but not BRCA1-proficient cells was seen previously in a screen setting by Adam *et al*. (Figure 2B) (1). The E3 ligase RNF8 functions upstream of 53BP1 and BRCA1 in DSB repair. Since loss of RNF8 results in impaired 53BP1 recruitment to DSBs (63,64), this might resemble 53BP1 loss which has previously been shown to reactivate HR in *BRCA1*-deficient cells (65,66), explaining the improved proliferation. On the other hand, RNF8 and RNF168 provide an alternative mode of PALB2 recruitment in a *BRCA1*-deficient setting and loss of RNF168 has been found lethal to cells lacking the recruitment mode of PALB2 through BRCA1 (67-69). Furthermore, loss of RNF8 in several BRCA1-mutant and -deficient cells promotes DNA damage through replication fork instability and R-loop accumulation (70). Hence, more research is needed to better understand the interplay between RNF8 and BRCA1 and how RNF8 deficiency mediates different phenotypes in *BRCA1*-deficient versus BRCA1^R1699Q^ mutated cells.

Importantly, loss of *PARP1* or *POLQ* led to reduced toxicity for BRCA1^R1699Q^ mutated cells compared to BRCA1-depleted cells. Previously, *POLQ* sensitivity in BRCA1-mutated cells was found to be dependent on functional end-resection in the targeted cells (71). This exemplifies that there might be differences in vulnerabilities for different BRCA1 mutations. *PARP1* is a well known target for treatment of tumours that carry BRCA1 mutations classified as pathogenic. BRCA1^R1699Q^ is marked as pathogenic in ClinVar (57) and hence these patients would qualify for PARPi treatment. Our data, however, provide a word of caution here as PARPi or POLQi might not be as effective to these tumours as expected. These differences between BRCA1-depleted and BRCA1^R1699Q^ cells highlight the importance to differentiate between patients with different BRCA1 mutations in the choice of treatment.

Individual mutations might even give rise to specific vulnerabilities, as we show here for BRCA1^R1699Q^ cells. In our hands, BRCA1^R1699Q^ cells, but not BRCA1-depleted cells, suffered from gross genomic instability in the form of increased micronuclei formation and anaphase bridges upon loss of the mitotic spindle associated protein NDE1. Previous research has shown that loss of either BRCA1 or NDE1 leads to an increase in DSBs and both proteins play a role in the regulation of centrosome duplication (3,36,58-60,72). Why there is specific synthetic lethality upon NDE1 loss in BRCA1^R1699Q^ cells remains elusive and requires more research. Perhaps the physical presence of BRCA1^R1699Q^, compared to full loss of the protein, results in differential signals to the cell making them more dependent on NDE1-mediated processes.

In conclusion, our data indicate the importance to study individual patient-derived mutations and not only focuss on full knockout of a gene when searching for novel therapeutic options. Furthermore, our data also indicate that comparing vulnerabilities between cells with different missense mutations and full knockouts presents a way to predict the severity of the functional consequences of a mutation.

## ACKNOWLEDGEMENTS

We thank the Luijsterburg lab (Leiden University Medical Center, The Netherlands) for providing CSA reagents, the Jonkers lab (Dutch Cancer Institute, The Netherlands) for providing HCC1937 cells and Maaike Vreeswijk (Leiden University Medical Center, The Netherlands) for critically reading our manuscript. This work was financially supported by Oncode Institute funds for SMN and a Dutch Research Council (NWO) VIDI grant (VI.Vidi.192.039) to SMN.

## AUTHOR CONTRIBUTIONS

The study was designed by SMN with input from AS. All experiments were performed by AS, with help of MMG, RAK, MSMA, IN and VG. CRISPR screens were performed by AS and KdL and analysed by KdL. RMFW supervised KdL and was involved in discussions on the progress of the project. The manuscript was written by AS and SMN with input from all co-authors.

## MATERIAL AND METHODS

### Cell lines

RPE1 hTERT *PAC^-/-^ TP53^-/-^* cells containing endogenously tagged BRCA1-GFP-mAID were described previously (3). To deplete AID-tagged BRCA1, cells were treated with 500 µM auxin (Sigma Aldrich, Saint Louis, MO, USA; stock solution 35 mg/mL in EtOH) for 48 hours, unless stated otherwise. Introducing the mAID-GFP-tag made the cells resistant to puromycin and therefore subsequent nucleofection of pLentiCRISPRv2 containing sg*PAC*: ACGCGCGTCGGGCTCGACAT was performed to resensitize the cells to puromycin. Cells were clonally expanded and genotyping was performed by PCR amplification and Sanger sequencing of the targeted locus, followed by TIDE analysis (73).

RPE1 hTERT *PAC^-/-^ TP53^-/-^ BRCA1-mAID* with doxycycline inducible (Tet-On) Cas9 were obtained by transduction of the cells with Edit-R Inducible Lentiviral hEF1a-Blast-Cas9, selection and clone selection. Thereafter the same inducible Cas9 clone was transduced with a vector containing a Tet-On promotor and either BRCA1 wildtype, BRCA1^I26A^ or BRCA1^R1699Q^. For each BRCA1 variant, clones were selected based on the expression of BRCA1 and the phenotype assessed by BRCA1 and RAD51 IRIF. RPE1 hTERT *PAC^-/-^ TP53^-/-^ BRCA1-mAID* with inducible Cas9 were transduced with pLentiGuide-GFP-sg*NDE1* (sg*NDE1*: GAGAGACATGATCGTGGCGC) to obtain *NDE1^-/-^* cells. Cells were treated with 2 µg/mL of puromycin for selection, subsequently clonally expanded, and genotyping was performed by PCR amplification and Sanger sequencing of the targeted locus, followed genotype analysis using Synthego ICE (Synthego Performance Analysis, ICE Analysis. 2019. v3.0. Synthego). Subsequently the *NDE1^-/-^*cells were complemented virally with either EV, Tet-On BRCA1 wildtype, or Tet-On BRCA1^R1699Q^.

RPE1 hTERT *PAC^-/-^ TP53^-/-^ BRCA1^-/-^* cells were generated by multiple nucleofections of pLentiCRISPRv2 containing either sg*PAC*: ACGCGCGTCGGGCTCGACAT; sg*TP53*: CAGAATGCAAGAAGCCCAGA; or sg*BRCA1*: AAGGGTAGCTGTTAGAAGGC. Subsequently, cells were clonally expanded, genotyping was performed by PCR amplification and Sanger sequencing of the gRNA targeted locus, followed by TIDE analysis (73). Thereafter the cells were transduced with a vector containing a Tet-On promotor and either empty vector, BRCA1 wildtype or BRCA1^R1699Q^.

HEK 293T cells were obtained from ATCC (Manassas, VA, USA). RPE1 and HEK 293T cells were cultured in Dulbecco’s Modified Eagle’s Medium (DMEM, high glucose, GlutaMAX™ and pyruvate supplemented (ThermoFisher Scientific, Waltham, MA, USA)) + 10% Fetal Calf Serum (FCS) and 1% penicillin + streptomycin (Pen-Strep).

HCC1937 cells (gift from Prof. J.M.M. Jonkers, Dutch Cancer Institue) were cultured with Roswell Park Memorial Institute (RPMI) 1640 medium (ThermoFisher Scientific) supplemented with 10% Fetal Calf Serum (FCS) and 1% Pen-Strep. HCC1937 cells were transduced with either pCW57.1_Zeo_BRCA1-Ty1 to rescue BRCA1 expression or with the empty pCW57.1 vector as a control.

All cells were maintained at 37°C, 5% CO_2_. When BRCA1-depleted, cells were maintained at 37°C, 5% CO_2_, 3% O_2_. All cells were regularly checked for mycoplasma contamination.

### Antibodies

The following primary antibodies were used for western blotting: Mouse α BRCA1 (Merck, OP92; 1:1,000), Mouse α Cas9 (Cell Signalling Technology, 7A9-3A3 #14697; 1:1,000), Rabbit α ERCC8/CSA (Abcam, ab137033; 1:500), Rabbit α NDE1 (ProteinTech, 10233-1-AP; 1:1,500), Mouse α TUBULIN (Sigma, T6199; 1:5,000), Mouse α TY1 (diagenode, C1520054, 1:1,000). The following secondary antibodies were used for western blotting: Goat α Mouse or Goat α Rabbit labelled with respectively IRDye 800 or IRDye 680 (LI-COR; 1:15,000) and HRP-labelled Goat α Mouse and Donkey α Rabbit (ThermoFisher Scientific, 1:5,000).

The following primary antibodies were used for immunofluorescence: Rabbit α BRCA1 (Millipore, 07-434; 1:1,000), Rabbit α RAD51 (Bio Academia, 70-001; 1:15,000), and Mouse α yH2AX (pSer 139, Millipore 05-636; 1:5,000). The following secondary antibodies were used for immunofluorescence: Goat α Mouse or Goat α Rabbit labelled with either Alexa 555 or Alexa 647 (Invitrogen; 1:1,000).

### Plasmids and cloning

All purification of plasmid DNA or PCR products was performed using commercially available kits (Qiagen, Hilden, Germany) according to the manufacturer’s protocol. Edit-R Inducible Lentiviral hEF1a-Blast-Cas9 Nuclease (CAS11229) to introduce doxycycline inducible Cas9 by transduction was obtained from Horizon (Cambridge, UK).

To obtain pCW57.1_Zeo_BRCA1-Ty1 or pCW57.1_Puro_BRCA1-Ty1, full length BRCA1 was PCR-amplified from pCL-MFG-BRCA1 (Addgene #12341 (74)) with a triple Ty1-tag included in the reverse primer and subsequently cloned into pENTR_1A. The BRCA1 mutations R1699Q and I26A were obtained by site-directed mutagenesis of BRCA1 in the pENTR_1A vector. Subsequently the BRCA1 coding sequence was transferred to pCW57.1_Zeo or pCW57.1_Puro containing a tetracycline-inducible promoter (Tet-On) using gateway cloning.

For the multi-colour competition assays, the sgRNAs targeting the described genes were cloned into the BsmBI-digested lentiviral expression construct pLentiGuide_GFP_puro (2). See Table 1 for used sgRNA sequences.

**Table 1.**
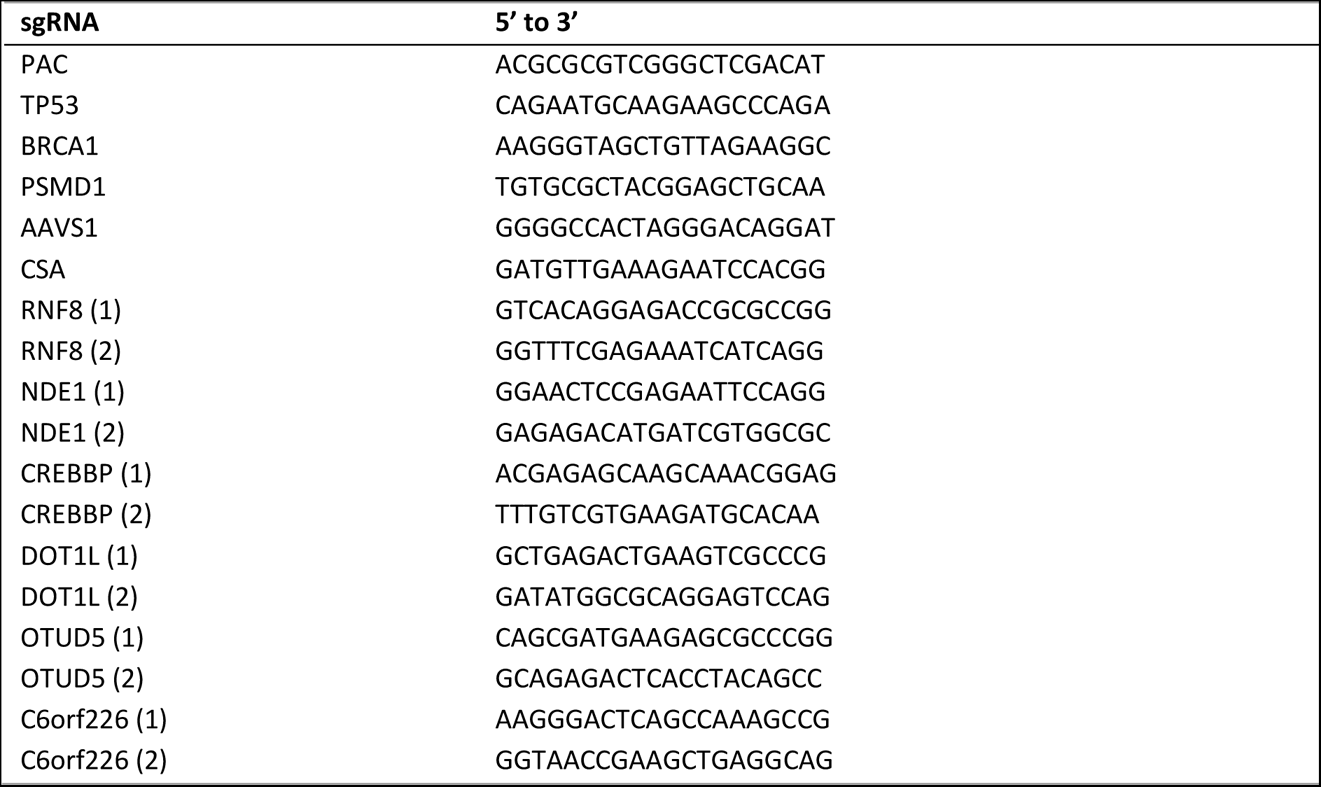
used sgRNAs throughout this study.

For the survival experiments the sgRNA targeting CSA (sg*CSA*: GATGTTGAAAGAATCCACGG) was inserted into pLenti-guide-OSS-mCherry by digestion of the vector with BsmBI (gift from B.A.F.J de Groot and Dr. M.S. Luijsterburg, Leiden University Medical Center).

### Viral transductions

Lentivirus was produced in HEK 293T cells by jetPEI transfection (Polyplus Transfection, Illkirch, France) of the Edit-R, pLentiGuide, pLenti-guide-OSS-mCherry, pLentiCRISPRv2, or pCW57.1 plasmid with third generation packaging vectors pMDLg/pRRE, pRSV-Rev and pMD2.G. Viral supernatants were harvested 48-72 hours post transfection, filtered (0.45 µm filter) and used to transduce cells at an MOI of ∼1 in the presence of 4 µg/mL polybrene. For RPE1 *PAC^-/-^* cells, 2 µg/mL of puromycin, 2 µg/mL blasticidin, or 200 µg/mL zeocin was used to select successfully transduced cells. For HCC1937, 8 µg/mL of polybrene was used for transduction and 1 µg/mL of puromycin was used to select successfully transduced cells.

### SDS-PAGE and Western Blot

Cells were lysed for 30 minutes on ice in RIPA lysis buffer (1% NP-40, 50 mM Tris-HCl pH 7.5, 150 mM NaCl, 0.1% SDS, 3 mM MgCl_2_, 0.5% sodium deoxycholate) supplemented with Complete Protease Inhibitor Cocktail EDTA-free (Sigma Aldrich) and 100 U/mL Benzonase (Sigma Aldrich). LDS sample buffer with DTT was added to the lysates, followed by denaturation at 95°C for 5 minutes. Proteins were separated by SDS-PAGE on 4-12 % gradient gels (ThermoFisher Scientific) and transferred to Amersham Protran premium 0.45 µm nitrocellulose membrane (GE Healthcare Life Sciences, Chicago, IL, USA). Membranes were blocked with 5% skimmed milk (Santa Cruz Biotechnology, Dallas, TX, USA) in 1x TBS or Blocking buffer for fluorescent WB (Rockland, Pottstown, PA, USA), and stained with primary and secondary antibodies. After secondary antibody-staining, the membranes were imaged on an Odyssey CLx scanner (LI-COR BioSciences, Milton, UK). Alternatively, when HRP-labelled secondary antibodies were used, the membranes were treated with the WesternBright ECL HRP Substrate kit (Advansta, San Jose, CA, USA) and imaged on an Amersham Imager 680 (Bioké, Leiden, The Netherlands) or an iBright 1500 (Invitrogen, Carlsbad, CA, USA).

### Immunoprecipitation

Cells were seeded in two 15 cm dishes per condition and grown to 90% confluency. Cells were harvested 1 hour post irradiation with 5 Gy and lysed rotating at 4°C for 45 minutes in 1 mL ice-cold NETT buffer (50 mM TRIS pH 7.6, 100 mM NaCl, 5 mM EDTA pH 8.0, 1x Complete Protease Inhibitor Cocktail EDTA-free, 0.5% Triton X-100, 7 mM MgCl_2_) and 500 U Benzonase. Lysates were centrifuged at 14,000 rpm at 4°C for 10 minutes, supernatant was collected and 40 µL input was saved.

Dynabeads protein-G (ThermoFisher Scientific) were washed 3 times with NETT buffer and mixed with α-Ty1 (Diagenode, Liege, Belgium), followed by incubated for 2.5 hours in this buffer. Afterwards, the Dynabeads protein-G plus α-Ty1 were washed twice and incubated with the pre-cleared lysates for 4 hours. Subsequently, the lysates were washed 5 times with NETT buffer. 2x LDS sample buffer with DTT was added to the beads, followed by denaturation at 95°C for 5 minutes, SDS-PAGE and Western blotting.

### Immunofluorescence

For BRCA1 and yH2AX IRIF, cells were grown on sterile 13 mm glass coverslips to 85% confluency and fixed 3 hours post irradiation with 10 Gy. Cells were pre-extracted with ice cold nuclear extraction (NuEx) buffer (20 mM Hepes pH 7.5, 20 mM NaCl, 5 mM MgCl_2_, 1 mM DTT, 0.5% NP-40 (IGEPAL CA-630, Sigma Aldrich), 1x Complete Protease Inhibitor Cocktail EDTA-free (Sigma Aldrich) for 12 minutes at 4°C and directly fixed with 2% (w/v) paraformaldehyde in PBS (20 minutes at room temperature) afterwards.

For RAD51 IRIF, cells were grown on sterile 13 mm glass coverslips to 85% confluency and fixed 3 hours post irradiation with 10 Gy. Cells were fixed and permeabilized with 1% (w/v) paraformaldehyde, 0.5% Triton X-100 in PBS for 20 minutes at room temperature, washed three times with PBS, further fixed and permeabilized with 1% (w/v) paraformaldehyde, 0.3% Triton X-100 in PBS for 20 minutes.

Subsequently, the fixed and permeabilized cells were washed three times with PBS and blocked with PBS+ (5 g/L BSA, 1.5 g/L glycine in PBS) for 30 minutes at room temperature. Blocked cells were incubated 1.5 hours with the primary antibody in PBS+, washed 4 times with PBS, and incubated 1 hour with DAPI 0.1 µg/mL (stock 100 µg/mL) and the secondary antibody in PBS+. All antibody incubations were performed at room temperature.

For micronuclei and anaphase bridge analysis, cells were grown on sterile 13 mm glass coverslips to 85% confluency and fixed after 48 hours of auxin treatment. After fixation the cells were permeabilized with 0.3% Triton X-100 in PBS, blocked with PBS+ and incubated with DAPI as described above.

After washing 4 times with PBS the coverslips were mounted using Aqua-Poly/mount (Polysciences, Warrington, PA, USA). Pictures were made using the Zeiss Axio Imager 2 fluorescent microscope at a 40x zoom. Foci of at least 100 cells per condition per replicate were quantified using the IRIF analysis 3.2 Plugin in ImageJ (75). For micronuclei and anaphase bridge analysis at least 100 cells per condition per replicate were quantified.

### Clonogenic survival assay

After 48 hours of transduction with pLenti plasmid containing either sg*CSA* or sg*AAVS1*, RPE1 hTERT cells were seeded in 10-cm dishes (RPE1 *BRCA1-mAID*: 500 cells) and treated as indicated. Medium containing Olaparib (16 nM or 50 nM) (Selleck Chemicals, Planegg, Germany), doxycycline (1 µg/mL) and/or auxin (500 µM) was refreshed after 7 days. After 14 days, colonies were stained with crystal violet (0.4 % (w/v) crystal violet, 20% methanol) and counted manually.

### Cas9 functionality test

Cells were transduced with pLentiGuide_GFP_puro_sgPSMD1 (targeting an essential gene, see Table 1 for used sgRNA sequences) or a control as described above and transduced cells were selected with puromycin. Cells were seeded at 10% confluency in a 6 wells plate, the next day medium was replaced and cells were grown in the presence or absence of doxycycline (1 µg/mL). The confluency was monitored during 7 days.

### Growth speed determination

0.5 million cells were seeded in a T75 flask and grown for 3 days. After 3 days the cells were counted. The doubling time was calculated: LN(2)/growth rate per day.

### Genome-wide CRISPR-Cas9 knockout screen and analysis

The screen was performed using clonally derived populations from RPE1 hTERT *PAC^-/-^ TP53^-/-^ BRCA1-mAID* cells with inducible Cas9 and transduced with either pCW57.1_Zeo_BRCA1^R1699Q^ or pCW57.1_Zeo_BRCA1^I26A^. Three populations of each cell line were transduced with the TKOv3 lentiviral library (gift from Prof. J. Moffat) in the presence of 8 µg/mL polybrene. Transduced cells were selected with 5 µg/mL puromycin for three days, after which t=0 samples were harvested for the three replicates per cell line and 20 × 10^6^ cells (corresponding to 280x coverage) were grown per replicate per condition, in the presence of doxycycline (1 µg/mL) to induce expression of Cas9 and BRCA1-variants. The four conditions in the screen were: **1)** *BRCA1-mAID* without auxin so BRCA1-proficient, **2)** *BRCA1-mAID* with 500 µM auxin treatment readily depleting BRCA1, **3)** *BRCA1-mAID* +BRCA1^R1699Q^ with auxin and and **4)** *BRCA1-mAID* +BRCA1^I26A^ with auxin. After 12 doublings, adapted for the growth speed of each specific cell line, each population was harvested and cells were stored at -80°C before DNA isolation.

The Blood and Cell Culture DNA Maxi kit (Qiagen) was used to isolate genomic DNA from each population at t=0 and end point. The DNA of 20 × 10^6^ cells was used as a template to amplify part of the lentiviral insert as described before in van der Weegen, de Lint *et al.* using the KAPA HiFi ReadyMix PCR kit (Roche, Basel, Switzerland) with TKO outer forward and reverse primers (76). Subsequently a second PCR assay was performed, attaching Illumina adapter sequences and containing a different Illumina i7 index sequence for each sample. Both the first and second PCR were checked on gel and the second PCR was purified using the QIAquick PCR Purification kit (Qiagen). Samples were mixed in equal amounts and sequenced on an Illumina NovaSeq 6000. Sequencing reads were mapped to the TKOv3 library sequences not allowing any mismatches, and the data were analysed using the software drugZ (v.1.1.0.2) as described before (76,77) or IsogenicZ, a DrugZ adaptation that includes a normalisation of end-point sgRNA counts based on the counts at t=0 (https://github.com/kdelint/IsogenicZ).

BioVenn (78) was used to create a visual representation of all the hits from each screen arm compared to the BRCA1-proficient arm of the screen. For the Volcano plot the fold change (Log2) was plotted against the significance (-Log10 FDR) of each individual gene.

STRING pathway analysis (79) was used for all hits (FDR< 0.1) that were synthetic lethal with BRCA1-depletion. Data for the analysis was based on textmining, experiments and databases. The FDR was corrected for multiple testing within each category using the Benjamini-Hochberg procedure. For the Panther pathway analysis, all genes with an normZ score of -2.5 or lower from the BRCA1-depleted (213 genes) or BRCA1^R1699Q^ (154 genes) arm were used as input for a statistical overrepresentation test of reactome pathways (version 85) using Panther version 18.0 (https://www.pantherdb.org). Pathways enriched with an FDR < 0.05 were considered statistically significant. The identified pathways per arm were manually curated for pathways involved in mitosis and compared between the two arms.

### MTT viability assay

After 48 hours of transduction with pLenti plasmid containing either sg*CSA* or sg*AAVS1*, HCC1937 cells were seeded in a 96 wells plate. After 6 days, the medium was removed from the cells and medium with 10% MTT solution was added to the cells. After incubation at 37°C for 2-4 hours the medium was removed. Freshly prepared extraction solution (1M HCl:Isopropanol, 1:25) was added to the wells and incubated at 37°C for 15 minutes. Plates were measured with SpectraMax iD3 (Molecular Devices), with absorbance at 570 nm.

### Multi colour competition assay

RPE1 hTERT PAC-/-TP53-/-BRCA1-mAID with inducible Cas9 were transduced with virus of pLentiGuide_GFP containing a sgRNA (see Table 1 for sequences) for the gene of interest or pLentiGuide_mCherry_LacZ_sgRNA. Transduced cells were selected by puromycin (2 µg/mL) treatment for 48h, and subsequently, GFP- and mCherry-positive cells were mixed 1:1. Mixed populations were seeded at 10,000 cells per well in a 12-well plate and treated with doxycycline (2 µg/mL) and auxin (500 µM). Cells were imaged every 4 days for a total of 20 days using an ArrayScan Cellomics high content microscope (Thermo Fisher Scientific). HCS Studio Cell Analysis Software (Thermo Fisher Scientific) was used to quantify the number of GFP- and mCherry-positive cells per well. Cells were passaged upon confluency. To assess gene editing efficiencies of the used sgRNAs, 4 days post treatment with 2 µg/mL doxycycline (7 days post transduction), genotyping was performed by PCR amplification and Sanger sequencing of the targeted locus, followed by analysis with Synthego ICE (Synthego Performance Analysis, ICE Analysis. 2019. v3.0. Synthego).

---

**Supplemental figure 1.**
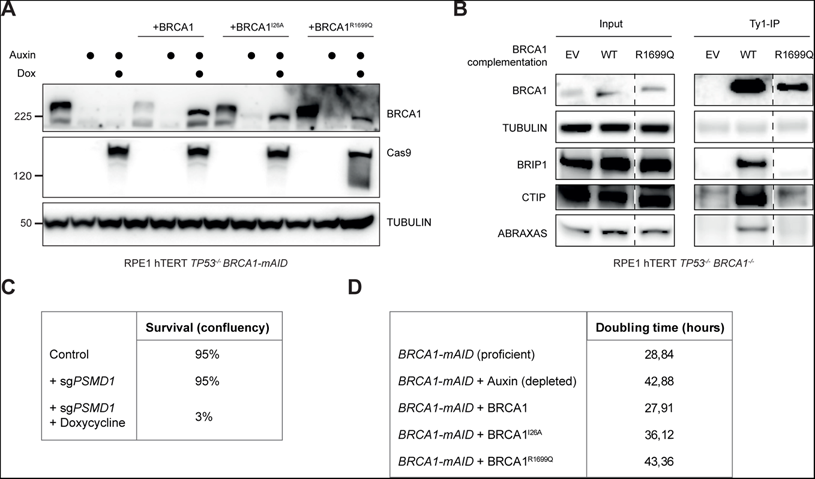
Genome-wide CRISPR-Cas9 knockout screen preparation. (A) The expression of BRCA1 and Cas9 was assessed by Western blotting of the indicated complemented RPE1 hTERT *TP53^-/-^ BRCA1-mAID* + doxycycline inducible Cas9 cells lines. Data shown represent three independent experiments. **(B)** Ty1 immunoprecipitation on RPE1 hTERT *TP53^-/-^ BRCA1^-/-^* cells complemented with BRCA1-Ty1 and BRCA1^R1699Q^-Ty1. Dashed line indicates removal of non-relevant lanes post-imaging. **(C)** The eficiency of virally integrated inducible Cas9 cassette was assessed by survival of the cells after transduction with a sgRNA against *PSMD1*, an essential gene. Data shown represent two independent replicates. **(D)** The growth speed of each indicated RPE1 hTERT *TP53^-/-^* cell line was assessed by monitoring cell growth and calculating the doubling time. Data shown is the average of two independent replicates.

**Supplemental figure 2.**
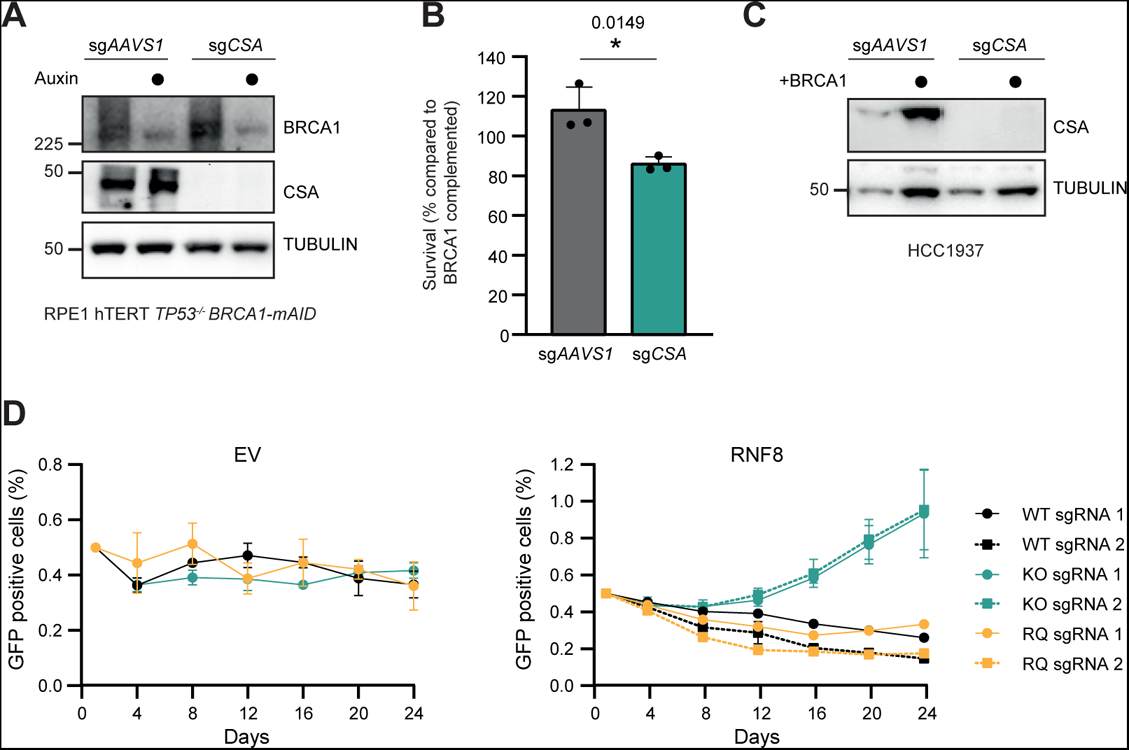
BRCA1-depleted cells are sensitive to CSA loss. (A) Lysates of the RPE1 hTERT *TP53*^-/-^ *BRCA1-mAID* cell lines described in Figure 2D were analysed by western blotting. **(B)** HCC1937 cell lines either compleme-mented with EV or *BRCA1* cDNA, were virally transduced to express *Cas9* cDNA and a sgRNA against *AAVS1* or *CSA*, followed by a MTT viability assay (n=3, mean+SD, *p<0.05, ratio paired t-test)). Western blot of lysates shown in Supple-mental figure 2C. **(C)** Lysates of the HCC1937 cell lines described in Supplemental figure 2B were analysed by western blotting. **(D)** RPE1 hTERT *TP53*^-/-^ *BRCA1-mAID* cells expressing doxycycline inducible Cas9, either BRCA1-proficient (WT, black lines), BRCA1-depleted with auxin (KO, green lines) or BRCA1^R1699Q^ complemented (RQ, orange lines), were infected with indicated *RNF8* sgRNA together with GFP, or with an empty vector together with mCherry. GFP- and mCherry-positive cells were mixed 1:1, and the ratio of GFP-positive cells in the population was determined over time (n=3, mean±SEM). Average eficiency of the sgRNA1 against *RNF8*, determined by using Synthego ICE (Synthego Performance Analysis, ICE Analysis. 2019. v3.0. Synthego) was as follows: 69% in the BRCA1-proficient (WT) cells (including BRCA1-depleted cells after auxin treatment; KO), and 73% in the BRCA1^R1699Q^ complemented cells (RQ).

**Supplemental figure 3.**
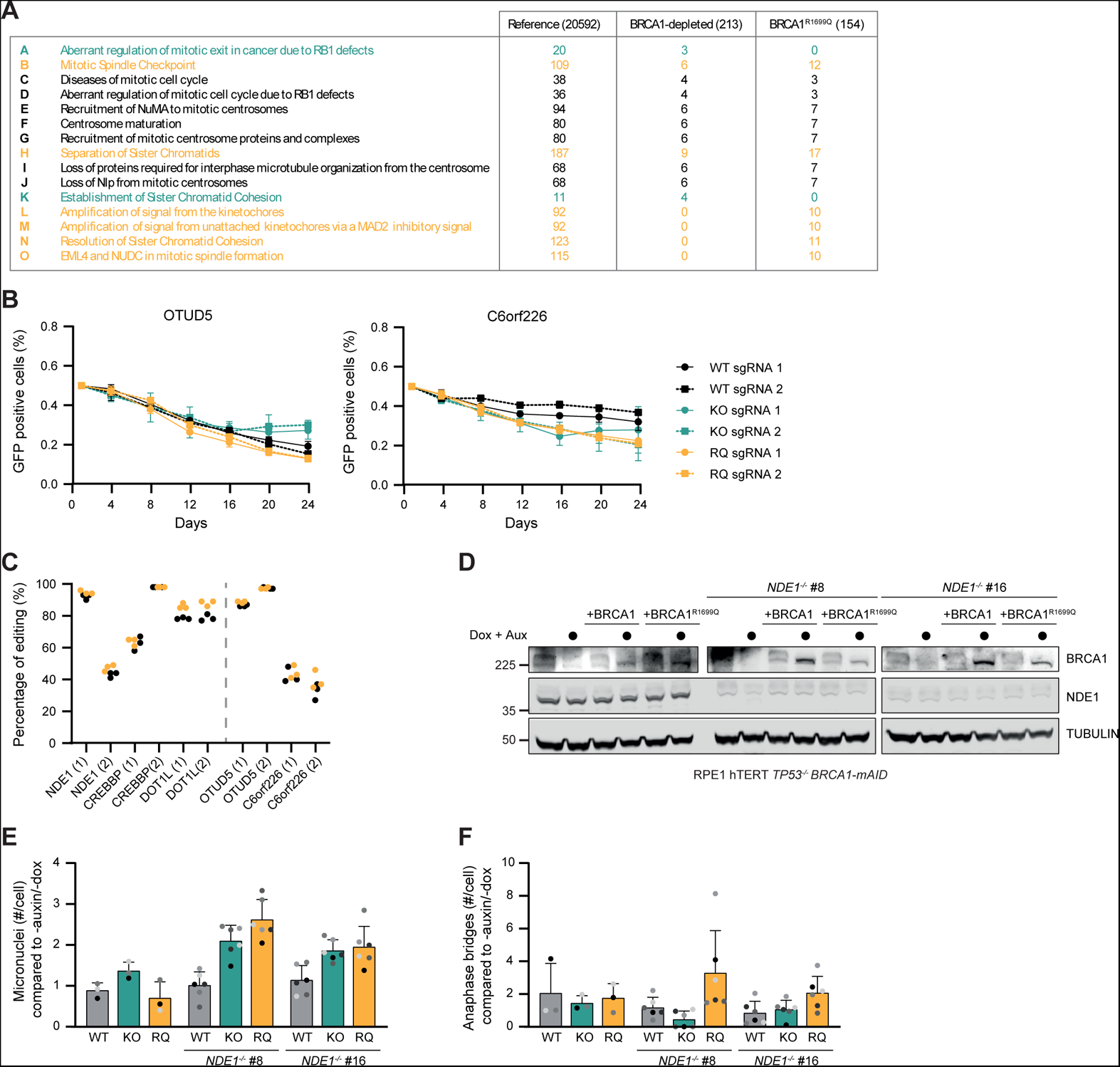
NDE1*-/-* cells were succesfully made and complemented. (A) Panther pathway analysis on the top vulnerabilities (NormZ < -2.5) of BRCA1-depleted cells and BRCA1^R1699Q^ cells. Table shows the raw number of genes involved in the specific mitotic process for the two different BRCA1 statuses as shown in Figure 4A. **(B)** RPE1 hTERT *TP53*^-/-^ *BRCA1-mAID* cells expressing doxycycline inducible Cas9, either BRCA1-proficient (WT, black lines), BRCA1-depleted with auxin (KO, green lines) or BRCA1^R1699Q^ complemented (RQ, orange lines), were infected with indicated sgRNA together with GFP, or with an empty vector together with mCherry. GFP- and mCherry-positive cells were mixed 1:1, and the ratio of GFP-positive cells in the population was determined over time (n=3, mean±SEM). gRNA efficiency is shown in Supplemental figure 3C. **(C)** Efficiency of each indicated sgRNA per replicate, determined using Synthego ICE (Synthego Performance Analysis, ICE Analysis. 2019. v3.0. Synthego). Black data points indicate the sgRNA efficiency in BRCA1-proficient (WT) cells (including BRCA1-depleted cells after auxin treatment; KO), orange data points in BRCA1^R1699Q^ complemented cells (RQ). **(D)** Western blot to check the expression of BRCA1 and NDE1 in the lysates decribed in Figure 4C and Supplemental figure 3E,F. **(E)** The indicated complemented (with either EV, doxycycline inducible BRCA1-WT or BRCA1^R1699Q^) RPE1 hTERT *TP53*^-/-^ *BRCA1-mAID NDE1^+/+^* or *NDE1^-/-^* cell lines were grown with or without auxin (to deplete endogenous BRCA1) and doxycycline (to express the BRCA1 complementation) for 48 hours before fixation. Micronuclei per cell were quantified (n=3 and n=6, mean+SD). **(F)** The indicated complemented (with either EV, doxycycline inducible BRCA1-WT or BRCA1^R1699Q^) RPE1 hTERT *TP53*^-/-^ *BRCA1-mAID NDE1^+/+^* or *NDE1^-/-^* cell lines were grown with or without auxin (to deplete endogenous BRCA1) and doxycycline (to express the BRCA1 complementation) for 48 hours before fixation. Samples were analysed for anaphase bridges (n=3 and n=6, mean+SD).

